# Sand-Mediated DNA Extraction From Orchids For Genomic Applications

**DOI:** 10.1101/2025.10.08.681153

**Authors:** Carlos Hernán Barrera-Rojas, Cássio van den Berg

**Affiliations:** Universidade Estadual de Feira de Santana (UEFS), Laboratório de Sistemática Molecular de Plantas (LAMOL), Av. Transnordestina, s.n., 44036-900, Feira de Santana, Bahia, Brazil

**Author notes:** Corresponding author: **Cássio van den Berg**, Universidade Estadual de Feira de Santana Departamento de Ciências Biológicas, Av. Transnordestina, s/n. CEP.: 44036-900 Feira de Santana-Bahia Brazil.

**Keywords:** DNA extraction, Orchids, Genetics, Molecular biology, CTAB methods

## Abstract

DNA extraction is an essential routine procedure for downstream applications in genetics and molecular biology. Since DNA was extracted for the first time, countless protocols have been developed yielding excellent results; however, the diversity of plant families and the distribution in remote areas represent a challenge for these protocols. Orchidaceae, one of the most species-rich plant families in the world, is a group with abundant polyphenols and polysaccharides in its tissues. In addition, sample collection in remote areas represents a challenge, consequently, conventional protocols can be inappropriate for obtaining DNA. In this way, CTAB-based gel preservation has emerged as an excellent alternative to easily collect, store and transport samples, moreover, alternatives such as sterile sand and DNA stabilization solutions during grinding allow a suitable DNA extraction. Here, we combined saturated NaCI-CTAB gels, sterile sand, sucrose-based DNA stabilization buffer, and conventional CTAB solution for obtaining DNA from fresh and stored samples for up to 32 days at room temperature, and described, from sample collection to spectrophotometer-obtained quantification, a non-organic method for obtaining DNA from orchids species for genomic applications. This protocol allows not only the extraction of suitable quantities of high molecular weight DNA of adequate purity and integrity from different orchid species without the use of expensive equipment and cumbersome reagents such as liquid nitrogen and toxic phenols, and/or time-consuming procedures, but also, the use of this DNA in downstream applications such as PCR-based genotyping, restriction enzyme analysis, and most genomic applications.

## INTRODUCTION

DNA extraction is a conventional and routine procedure in molecular biology laboratories for downstream applications such PCR-derived analysis (Meneghetti et al., 2014), high-resolution melting genotyping (Kim et al., 2023), barcoding for species identification (Bhargava and Sharma, 2013), NGS-based phylogenetic analysis (Draper et al., 2022), plant pathogen detection (Nezhad, 2014), and digestion by restriction endonucleases (Smith, 1993), among others. This huge spectrum of applications makes DNA extraction a key factor for obtaining excellent results in high-throughput applications. In general, there are two routes for DNA extraction: physical and chemical methods (Gupta, 2019). Physical extraction includes magnetic beads and paper-based methods, and chemical extractions involve organic and non-organic compounds (Sheershika and Ram, 2024). Organic DNA extraction depends mainly on phenol-chloroform and isoamyl alcohol (Thomas et al., 1989), and the non-organic extraction relies on Proteinase K, Sodium Dodecyl Sulfate (SDS), salting out, silica-gel-based techniques and Cetyltrimethylammonium bromide (CTAB) method (Gupta, 2019; Rivero et al., 2006; Janson and Tischler, 2012; Clarke, 2009; Connon, 2023; Lee et al., 2021; Doyle and Doyle, 1987; Doyle, 1991; Saghai-Maroof et al., 1984)This non-organic methodology can be made using commercially available kits and normally are dependent on liquid nitrogen (LN); nevertheless, commercial kits-based DNA extraction, offered by several companies, and LN-dependent tissue grinding can be expensive, especially with high amounts of samples and inaccessible in remote areas.

Conventional DNA extraction involves several steps, including sample collection, cell wall breaking, cell membrane disruption, and finally DNA precipitation and elution. Depending on the purpose, sample collection can be normally collected into 1.5 mL microcentrifuge tubes and immediately frozen into LN (Maier et al., 2010), one of the most common alternatives for laboratories; however, it is not available in several cases, especially in remote locations, due to transportation and evaporation rates of LN. Previous reports have demonstrated that DNA suitable for downstream analysis can be obtained from leaves that have been stored in CTAB-based gels for a short period of time at ambient temperature and for over one year at -20°C (Rogstad, 1992). Therefore, saturated NaCI-CTAB gels become an excellent alternative for collecting and storing samples because the ingredients are readily available and easy to transport.

In order to release cellular components, DNA extraction must involve breaking down the cell wall and disrupting the cell membrane. Cell wall rupture can be obtained by grinding the tissue using dry ice or liquid nitrogen, nevertheless, it has been reported that sterile sand provides a means of grinding the samples providing DNA yield suitable for any application (Arif et al., 2010). In addition, Sucrose-Tris-EDTA (STE) buffer has been used as helpful for DNA stabilization, during grinding, protecting the samples from degradation (Takakura and Nishio, 2012). Cell membrane disruption can be obtained by using a detergent solution, normally sodium dodecyl sulfate (SDS) or CTAB. The SDS is a powerful detergent that breaks down cellular membranes and denatures proteins, by disrupting non-covalent bonds, and denaturing them (Janson and Tischler, 2012). CTAB is a surfactant that removes membrane lipids and promotes cell lysis; also, when the tissue contains high amounts of polysaccharides, CTAB binds and removes them from solution (Clarke, 2009); several DNA extraction protocols using SDS our CTAB are available (De Silva et al., 2025; Russo et al., 2022; Sahu et al., 2012; Emaus et al., 2022; Clarke, 2009; Porebski et al., 1997; Doyle and Doyle, 1987; Doyle, 1991). Combining these methods, storage samples in CTAB-based gels, grinding with sterile sand and STE buffer, and CTAB-mediated cell lysis result in an excellent low-cost alternative with high DNA yield and purity, especially for plant families with samples from remote locations such as Orchidaceae.

Orchidaceae is one of the most species-rich plant families with about 28,000 currently accepted species representing about 10% of the world’s flowering plants (Roberts and Dixon, 2008). This group of plants has a diverse lifestyle and grows as epiphytes, terrestrials, and rupicolous in different ecosystems (Zhang et al., 2017), and due to its enormous number of species and wide geographical distribution, this family has been divided, based on phylogenetic studies, into 5 sub-families: Apostasioideae, Vanilloideae, Cypripedioideae, Epidendroideae, and Orchidoideae that are distributed worldwide specially in the tropics (Christenhusz and Byng, 2016). Studies of the Orchidaceae family mainly involve morphological and molecular-analysis based phylogeny. These analyses rely strictly on the extraction and purification of high-quality nucleic acids. The DNA extraction from orchids has been previously reported from fresh tissues or dried herbarium specimens with results that allowed obtaining suitable DNA for those analyses. (Kamba and Ranjan, 2018; Lim et al., 1998; Mazo et al., 2012); however, DNA extraction from CTAB gels-preserved samples from orchids has not been reported; in addition, DNA extraction studies used unnecessary reagents, times, temperatures, and procedures making these protocols expensive, time consuming, and unsafe. Here, we have modified step-by-step the procedure for DNA isolation of fresh and preserved samples in CTAB gels in order to develop an updated methodology, maintaining the yield and purity of DNA for downstream applications. Finally, we outline step-by-step this process in order to offer a simple, safe, inexpensive, and rapid method of DNA extraction from orchids.

## PROTOCOL

In the following steps, we will describe the updated protocol for DNA extraction from orchids. In this study we have used plants from the Plant Molecular Systematics Laboratory at the State University of Feira de Santana (Brazil) that are growing under greenhouse conditions. With the exception of the family Apostasioideae, available exclusively in Australia, we have used species from the four orchid subfamilies: Vanilloideae, Cypripedioideae, Epidendroideae, and Orchidoideae. These plants were previously obtained from the natural environments with ICMBio authorization (SISBIO license 87099-1), which allows the collection of materials from all Orchidaceae species, including endangered species, from natural environments and National Parks from Brazil. In order to facilitate carrying out the experiments, we initially standardized the protocol for DNA extraction from orchids by initially using the species *Vanilla palmarum* Lindl. (Vanilloideae; Figure 1 A). Later, in order to validate the protocol, we have used species from the subfamilies previously mentioned (Figure 2).

**Figure 1.**
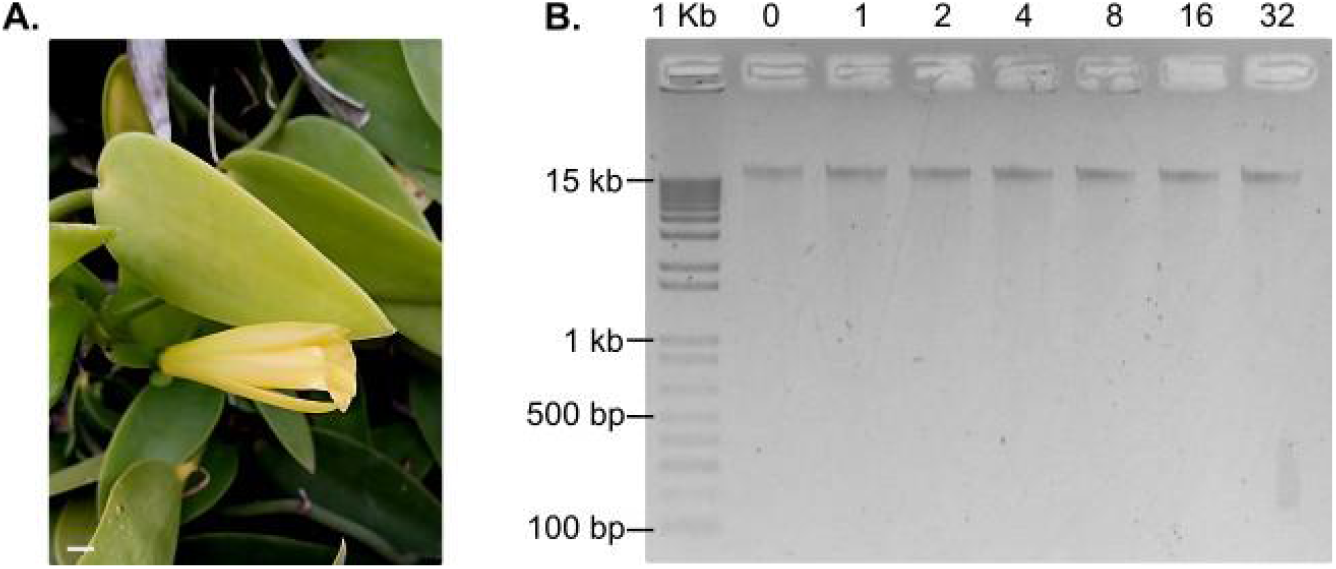
DNA extraction from samples of *Vanilla palmarum* (Vanilloideae) stored in NaCl-CTAB gel. **A**: picture of a Vanilla plant. Scale bar: 1 cm. **B**.: gel of extracted DNA from samples stored for 0, 1, 2, 4, 8, 16 and 32 days in NaCl-CTAB gel. Lane 1: molecular marker 1 Kb Plus DNA Ladder (ThermoFisher). Lane 2-8: 0, 1, 2, 4, 8, 16 and 32 days of storing.

**Figure 2.**
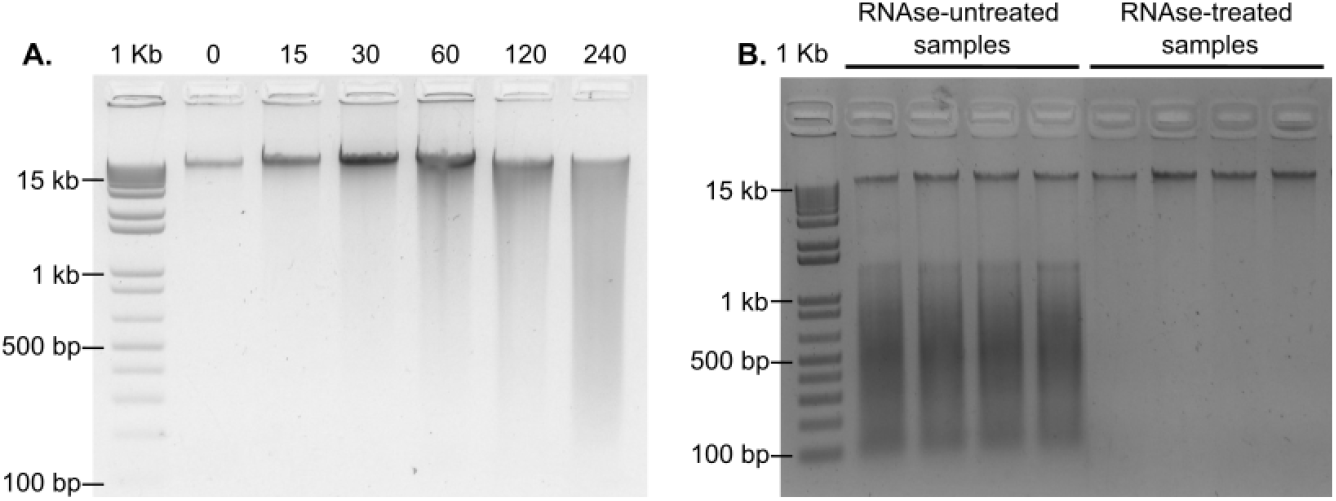
DNA extraction under heat incubation and RNAse treatment. **A**: Gel of extracted DNA from heat-treated *Vanilla* samples. Time incubation between 0 and 240 minutes. Lane 1: molecular marker 1 Kb Plus DNA Ladder (ThermoFisher). Lane 2-7: 0-240 minutes of incubation time respectively. **B**.: gel of extracted DNA from RNAse untreated and treated samples. 1% Agarose gels staining with Unisafe DYE (UniScience) run at 150 volts and 150 milliamps for 30 minutes.

### 1. Sample collection

Under laboratory conditions, it is usual to collect fresh samples and immediately grind them, nevertheless, this procedure is not always possible specially when collecting is carried out in remote areas; in addition, collecting, storing, and preserving samples under refrigerated conditions can be difficult and it is not possible for all plant taxa. To overcome this difficulty, saturated NaCl-CTAB gels have emerged to avoid DNA degradation (Rogstad, 1992). It is easy to prepare and transport, inexpensive, lasting and safe. Here, we have used a saturated NaCl-CTAB gel (35% NaCl + 3% CTAB). We recommend preparing and aliquoting 1.0-1.5mL in 2 mL free-DNAse microcentrifuge tubes, however it can be aliquoted, if required, in larger tubes such as 15 or 50 mL DNAse-free tubes. Also, CTAB gel can be stored indefinitely and transported at room temperature (Figure 3 A).

**Figure 3.**
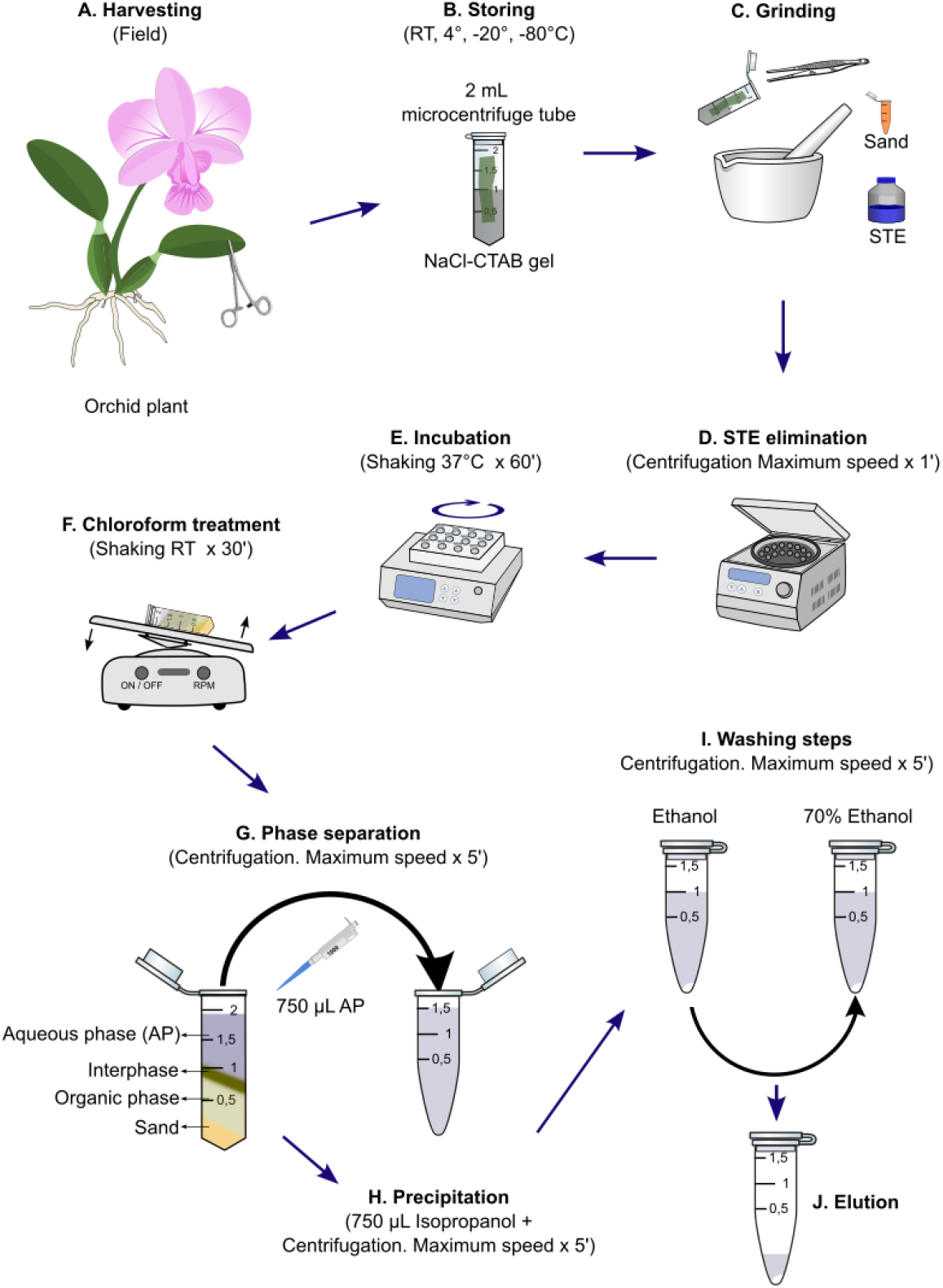
Flowchart of the DNA extraction process in orchids. Samples are collected (A) on NaCl-CTAB gels to be stored (B). After that, samples are grinded using STE and sand (C). The STE must be eliminated (D) to samples being incubated in CTAB (E) and chloroform treatment (F). Phase separation (G) and precipitation (H) and washing steps (I) are made by centrifugation. Finally, DNA elution (J).

### 2. Sample storage

DNA extraction is normally done immediately after collection and, depending on the purpose, samples can be stored in ultracold freezers at -80°C, however, this alternative is not applicable for samples that are collected in the field, such as orchid species. Thus, saturated NaCI-CTAB gels have emerged as an alternative to preserve DNA integrity (Rogstad, 1992). Orchid sample collection can be done at remote locations and samples must be stored for some time. However, information about time storage in high-salt CTAB gels is unknown. In order to evaluate if storing samples in saturated-salt CTAB gels preserves the DNA integrity over time, we have collected and stored the samples of *V. palmarum* (Vanilloideae; Figure 1 A) in NaCI-CTAB gels for 0, 1, 2, 4, 8, 16 and 32 days at room temperature (24°C +/-1°) under darkness condition. As observed in Figure 1 B, DNA integrity remains intact at all times, indicating that, until 32 days after collecting the samples, the NaCl-CTAB gel preserves DNA integrity at room temperature. Therefore, we recommend storing samples for up to 32 days in saturated NaCI-CTAB gels at room temperature in the dark (Figure 3 B).

### 3. Sample grinding (Cell wall broken)

After collecting, samples must be properly ground to extract DNA. This step is usually done by using dry ice, LN or directly on DNA extraction buffers. However, dry ice or LN can be expensive and not available depending on the laboratory location; in addition, DNA extraction buffers need to preserve the DNA integrity. Therefore, sterile sand together with sucrose-based DNA stabilization buffer is a viable resource for grinding the orchid tissues while preserving the DNA (Takakura and Nishio, 2012; Arif et al., 2010). Here, we have used autoclaved and dry sand, previously washed with 2% sodium hypochlorite and distilled water, as a grinder agent, and cold STE (8,5% Sucrose, 30 mM Tris, 200 mM EDTA) as a DNA preservation agent. We recommend preparing a stock solution of STE and maintaining it up to several months at 4°C; also, use +/-0.5 g of sand and 1 mL cold STE for 100 to 200 mg of tissue during sample grinding (Figure 3 C).

### 4. Cell membrane disruption

CTAB promotes cell lysis as well as removes membrane lipids, polysaccharides and polyphenolic compounds as previously shown (Clarke, 2009; Porebski et al., 1997; Sahu et al., 2012; Emaus et al., 2022; Doyle, 1991). Also, conventional methods for DNA extraction required an incubation time at 60°C. Because orchid plants have fleshy leaves with high levels of polysaccharides and polyphenolic compounds, CTAB constitutes an excellent reagent for extracting DNA from these species, however the time incubation required to proper DNA extraction is unknown. Thus, in order to define the appropriate time to incubate orchids samples in CTAB, we have incubated samples from Vanilla leaves for 0 min, 15 min, 30 min, 60 min, 120 min, and 240 min in 800 μL of extraction buffer (1.4 M NaCl, 100 mM Tris HCl pH 8.0, 20 mM EDTA, 2% Polyvinylpyrrolidone (PVP), and 2% CTAB) at 60°C under constant agitation at 150 RPM. As observed in figure 2 A, there are no significant differences in DNA integrity between 0 and 30 minutes of shaking; however, after 60 min of agitation, DNA samples begin to degrade, indicating that the longer the time, the greater the degradation of DNA. Here, we recommend no incubation at 60°C as previously described.

### 5. RNAse treatment

Besides polyphenolic compounds and polysaccharides, RNA is also a main contaminant in extracted DNA that can cause difficulties conducting downstream experiments (Arif et al., 2010); thus, RNAse treatment is recommended for removal of unwanted RNA. Several protocols recommend adding RNAse at the end of the extraction process, however, it can be done by adding the RNAse during the incubation time of the extraction buffer. By doing this in this step, both the degraded RNA and the RNAse will be eliminated in subsequent steps. Here, we use 1μL100 mg/mL RNAse A for removing all RNA during the buffer lysis step and incubate the samples at 37°C during 60 mins under constant shaking (Figure 2 B; Figure 3 E). If there is no shaker, it is possible to incubate the samples in a water bath, which also offers excellent results.

### 6. Chloroform treatment

Chloroform is an agent in purifying DNA by promoting phase separation, protein denaturation and increased DNA purity. Chloroform separates, by centrifugation, the aqueous and organic phases helping to isolate DNA from proteins and other contaminants; also, it denatures and precipitates proteins that may be present in the sample. Finally, by removing contaminants, chloroform helps enhance the purity of the DNA. Therefore, chloroform treatment is critical during DNA extraction. Many protocols combine chloroform and isoamyl alcohol (IAA). The IAA is several times more expensive than chloroform that acts as an anti-foaming agent, therefore, we do not use alcohol IAA in this step and have got excellent results. However, depending on the procedure, the most common ratio for the chloroform:isoamyl alcohol solution is 24:1. Here we add 800 μL of chloroform to the samples and shake them for 30 mins at room temperature.

### 7. Phase separation

Chloroform denatures and precipitates proteins and, by centrifugation, separates the aqueous phase, containing the DNA, from the organic phase, containing proteins and other cellular components. Thus, it is critical to centrifuge the samples at maximum speed. Conventional protocols recommend centrifugation during 10 mins at 4°C. Here we have used 5 mins of centrifugation at room temperature and get excellent results.

### 8. DNA precipitation

After phase separation, it is critical to carefully get the aqueous phase and transfer it into a new microcentrifuge tube to precipitate DNA by reducing the solubility of DNA, causing it to aggregate and, in most cases, become visible. Although isopropanol is used to precipitate DNA from the aqueous solution in most cases other alcohols such as methanol or ethanol can be used; however, methanol has a significant risk level due to its toxicity and side effects, and ethanol is used preferably for subsequent washes of DNA. Here we use isopropanol with excellent results. In the fume hood, we remove 750 μL of the aqueous phase (supernatant) into a new DNAse-free 1.5 mL tube, add 750 μL of ice-cold isopropanol to each sample, shake tubes by inversion until completely mixed, and directly centrifuge at maximum speed for 5 mins. However, for better results in DNA concentration, previous centrifugation, it is possible to store samples at -20°C for indefinite time and/or increase centrifugation time. This is also a safe stop point if delaying other stages of the protocol for later.

### 9. Ethanol washes

Once the DNA has been precipitated, it is washed in two steps using cold ethanol and cold 70% ethanol. Ethanol is used to eliminate the residual organic components such as chloroform or other components, and 70% ethanol is used to eliminate residual salts. Here, we use 1 mL in each step followed by 5 minutes of centrifugation at maximum speed.

### 10. Drying and eluting DNA

After ethanol washes, precipitated DNA must be air dry. Here, we normally dry the DNA for a maximum of 15 minutes at room temperature (+/-24°C). Longer times should be avoided to prevent excessive DNA dehydration. Then we elute the DNA pellet with 30 μL of DNAse-free water, however, it can be eluted in 1x TE buffer (10 mM Tris-HCl pH 8.0, 1 mM EDTA, pH 8.0), and stored at 4 °C for short-term use, or -20°C to -80°C for long-term storage. Although EDTA can interfere in Magnesium (Mg^2+^)-dependent downstream reactions, Low TE or TE Low EDTA (10 mM Tris-HCl pH 8.0, 0.1 mM EDTA, pH 8.0) can avoid interference in these types of reactions while preserving DNA.

## DISCUSSION

In molecular biology and genetics, DNA of adequate purity and integrity is a key factor for downstream application such as PCR analysis, barcoding, genotyping, sequencing, among others (Meneghetti et al., 2014; Kim et al., 2023; Bhargava and Sharma, 2013; Draper et al., 2022). For these purposes, DNA extraction is a conventional and routine method in molecular and genetics laboratories. Since CTAB-mediated DNA extraction was reported for the first time in plants (Doyle, 1991), countless protocols have been developed; however, plants vary greatly in their chemical compositions and obtaining pure high-molecular-weight DNA is difficult in some species making this routine method a challenge. Orchid species have high levels of polysaccharides and also polyphenolic compounds, thus, CTAB-mediated DNA extraction is an ally in molecular biology procedures; nevertheless, unnecessary use of reagents, times and procedures make this outdated, expensive, hazardous and laborious. Here, we show and update methodology for DNA extraction of orchis, from harvesting to quantification to make this methodology simpler, cheaper, faster, and safer. In addition, we recommend collecting young undamaged plant leaves because young leaves have a higher density of cells per volume and should yield more DNA (Funk et al., 2017); in addition, newly opened flower tissues also have the same qualities.

A proper DNA extraction involves several steps, including sample collection and storage, cell wall breaking and cell membrane disruption, RNAse and chloroform treatment, phase separation, DNA precipitation, washing and eluting. While sample collection can be done under laboratory conditions, in most cases it must be done in remote areas. Therefore, saturated NaCI-CTAB gels have become an essential tool for collecting, transporting, and storing samples in a practical, economical, and safe manner. Here, we have demonstrated that NaCI-CTAB gels preserve DNA integrity from orchid samples for up to 32 days at room temperature in dark conditions, helping to simplify DNA extraction. In addition, samples can be stored at 4°, -20°, and even down to -80°C; however, further data deserves to be done. During cell wall broken, most of the protocols use LN, however, sucrose-based DNA stabilization buffer such as STE together with sterile sand becomes a great tool for obtaining obtain high-yield and high-purity DNA suitable for any further application as previously shown (Takakura and Nishio, 2012; Arif et al., 2010) avoiding high costs and laborious logistics.

CTAB is a surfactant useful for isolation of DNA especially from tissues containing high amounts of polysaccharides such as orchid species (Clarke, 2009). Conventional protocols use CTAB at high temperatures, +/-60°C, leading to excellent results (Clarke, 2009; Porebski et al., 1997; Sahu et al., 2012; Emaus et al., 2022; Doyle, 1991; Doyle and Doyle, 1987; De Silva et al., 2025; Saghai-Maroof et al., 1984) however, in some cases it is necessary to evaluate the effect of this temperature in DNA integrity. Here, we have shown that heating samples at 60°C is unnecessary and also harmful to the integrity of the DNA of the samples after 30 minutes. Instead of heating, combining lysis buffer and RNAse treatment in the same step not only preserve DNA but also save time. In addition, most protocols use β-mercaptoethanol which prevents oxidative damage to DNA (Gerstein, 2001); although, it is weakly toxic and is an irritant to skin and eyes (Knight, 2001), here we have not used this reagent in our analyses and we have not had any problems either during the extraction or in further analyses, suggesting that DNA extraction may be done without β-mercaptoethanol reducing risks and costs. This is also related to the storage of the samples in the CTAB-NaCl gels, which practically eliminates tissue browning during the collecting stage, and allows dismissing the use of β-mercaptoethanol and its risks. Routine DNA precipitation and washing following the sequence isopropanol, ethanol, and 70% ethanol were maintained in this work. Also, we recommend drying the DNA pellet at room temperature for 15 mins and eluting DNA with DNAse-free water, low TE or TE low EDTA have shown excellent results in downstream applications.

## ACKNOWLEDGMENTS

We thank the National Council for Scientific and Technological Development (CNPq) from Brazil for the funding (Post-doctoral fellowships No 150676/2024-7).

## AUTHOR CONTRIBUTIONS

Conceptualisation: C.H.B.R.; Experiments: C.H.B.R.; writing: C.H.B.R.; review: C.H.B.R., and C. v d B. All authors have read and agreed to the published version of the manuscript.

